# Characterisation and development of aspirin inducible biosensors in *E. coli* Nissle 1917 and SimCells

**DOI:** 10.1101/414169

**Authors:** Jack Xiaoyu Chen, Harrison Steel, Yin-Hu Wu, Yun Wang, Jiabao Xu, Cordelia P. N. Rampley, Ian P. Thompson, Antonis Papachristodoulou, Wei E. Huang

**Affiliations:** Department of Engineering Science, University of Oxford, Parks Road, Oxford, OX1 3PJ, United Kingdom; Environmental Simulation and Pollution Control State Key Joint Laboratory, State Environmental Protection Key Laboratory of Microorganism Application and Risk Control (SMARC), School of Environment, Tsinghua University, Beijing100084, PR China

**Author notes:** corresponding author: Wei E. Huang Department of Engineering Science, University of Oxford, Parks Road, OX1 3PJ, Oxford, United Kingdom. The authors wish it to be known that, in their opinion, the first 3 authors should be regarded as joint First Authors.

**Keywords:** synthetic biology, aspirin/salicylate acid transcriptional regulation, biosensor, SimCells, *E. coli* Nissle 1917, SalR competitive binding

## Abstract

A simple aspirin-inducible system has been developed by employing the P_sal_ promoter and SalR regulation system originally from *Acinetobacter baylyi* ADP1, which has been cloned into *E. coli* for characterisation of gene circuits and induction of novel SimCells (simple cells). Mutagenesis at the DNA binding domain (DBD) and chemical recognition domain (CRD) of the SalR protein in *A. baylyi* ADP1 suggests that inactive SalR_i_ can compete with activated SalR_a_, occupying the binding position of P_sal_ promoter. The induction of the P_sal_ promoter was compared in two different designs in *E. coli*: simple regulation (SRS) and positive autoregulated system (PAR). Both regulatory systems were induced in a dose-dependent manner in the presence of aspirin in the range of 0.05-10 μM. Over-expression of SalR in the SRS system reduces both baseline leakiness and inducible strength of P_sal_ promoter. A weak SalR expression significantly improve the inducible strength, which is in a good agreement of the proposed hypothesis of SalR_i_/SalR_a_ competitive binding. The PAR system provides a feedback loop that fine-tunes the level of SalR, displaying inducible strength. A mathematical model based on SalR_i_/SalR_a_ competitive binding hypothesis was developed, which not only reproduces the observed experimental results but also predict the performance of a new gene circuit design. The aspirin-inducible systems were also functional in probiotic strain *E.coli* Nissle 1917 (EcN) and SimCells produced from *E. coli* MC1000 ΔminD. The well-characterised and modularised aspirin-inducible gene circuits would be useful biobricks for bacterial therapy in environment and medical applications.

## Introduction

Synthetic biology has the potential to engineer bacteria for disease diagnosis (1) and cancer therapy (2). Since bacteria colonise human skin, gastrointestinal tracts, the respiratory and reproductive systems (3), and many bacteria preferentially associate with tumours (2), they are ideal agents for diagnosis and therapy. Bacterial therapy has shown great potential in biomedicine, with applications including the regulation of a host organism’s energy metabolism (4), the delivery of drugs (5), as well as modulating chemotherapy, radiotherapy, and immunotherapy treatments for cancer (6).

Advances in the utilisation of feedback-based synthetic circuits for biomass yield optimization (7), spatial control of tissue regeneration (8), and bacterial diagnosis therapy (9-12) have been characterised and implemented in *in vivo* studies. An ideal bacterial design for environment and medicine should have the following traits. The specific bacterial chassis should be safe for human applications. The inducer should have no side effects on human health, and should be able to trigger different levels of gene expressions in response to different inducer concentrations. Gene circuits should have minimal cross-talk and only be triggered by a specific inducer, rather than molecules present naturally in the human body or in common diet. Finally, to achieve effective bacterial therapy the release of drugs (especially cytotoxic drugs for treatment of cancer) from engineered bacteria must be accurately and reliably controllable with tightly regulated promoters.

Aspirin is a safe and specific inducer, which has been used widely as an analgesic and anti-inflammatory drug since 1897 (13), and is as such ideal for human applications. The clinical safe dosage of aspirin for an adult is 75-300 mg per day, and therapeutic blood concentration for adults is between 111 and 555 μM, whereas toxic concentration is between 832 and 1665 μM (14). The biological half-life of aspirin is 2-3 h for low dosages and 15-30 h for large dosages (15). Salicylate (SA) and aspirin regulatory module nahR/Psal::xylS2 is *Psedomonas* derived gene circuit and it has been applied in regulated expression of *Salmonella spp.* genes at millimolar level (9).

In this study, a simple and sensitive aspirin/salicylate regulated P_sal_-salR system in *Acinetobacter baylyi* ADP1 has been characterised (16-18). As a member of the LysR-type transcriptional regulator (LTTR) superfamily, SalR controls the salicylate degradation pathways in *A. baylyi* ADP1 (16) and is the activator of its own promoter P_sal_ (16, 18). It has been shown that SalR regulation can be activated by aspirin in *A. baylyi* (18). We examined the P_sal_-SalR regulation mechanism and characterised this system’s modularity, and to demonstrate that this system functions when it is transplanted into a different chassis or network architecture. Furthermore, we proposed an inactive and activated SalR_i_/SalR_a_ competitive binding hypothesis and developed a novel mathematic model to describe the process. We designed three sensitive aspirin responsible P_sal_-SalR gene circuits including a positive autoregulated system (PAR) and two simple regulation system (SRS) with varying promoter strengths. The performance of P_sal_-SalR gene circuits has been characterised in *E. coli* DH5α, probiotic *E. coli* nissle 1917 (EcN) (19) and chromosome-free SimCells (20). A novel mathematical model not only quantitatively reproduced the experimental results but also predicted the performance of a new gene circuit design. The sensitive aspirin gene circuits were functional in both EcN and SimCells, which provide aspirin-controllable and safe tools for potential bacterial therapy in environment and medicine applications.

## Materials and Methods

### Chemical and reagents

All reagents were purchased from Sigma Aldrich (Dorset, UK) unless otherwise noted. Antibiotics (Kanamycin and Ampicillin) were obtained from Fisher Scientific (UK). Sterile 1X phosphate buffered saline (PBS) solution was made by diluting 10X PBS solution stock purchased from Fisher Scientific (UK).

### Strains and plasmids

The strains and plasmids used in this study are listed in Table 1.

**Table 1.**
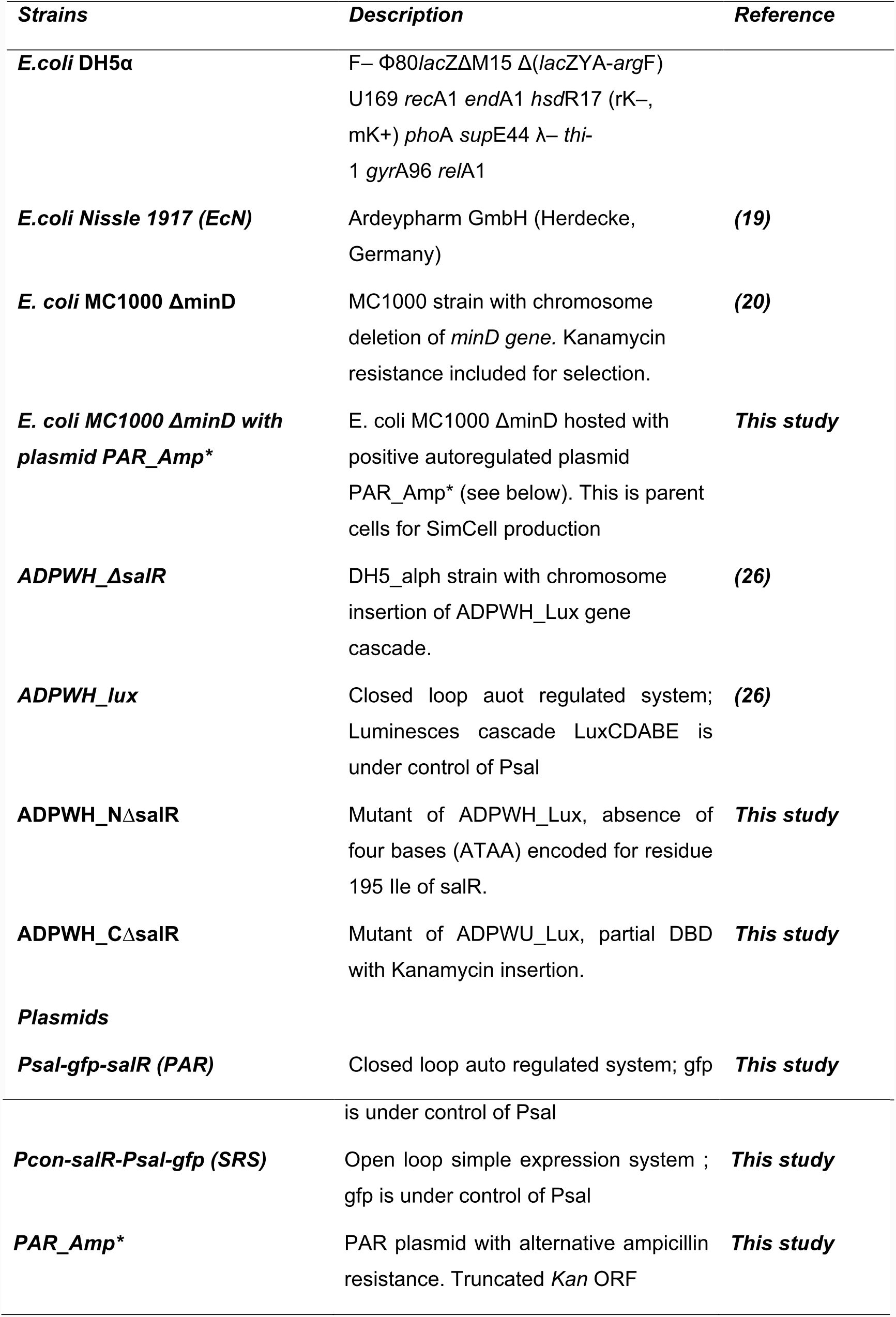
Bacterial strains and plasmids used in this study.

Plasmids (Table 1) used in this study were synthesized by GeneArts (UK) and transformed into *E.coli* DH5α cells which were made chemically competent beforehand. *E.coli* Nissle 1917 (EcN) was acquired from Ardeypharm GmbH(Herdecke, Germany). *E. coli* MC1000 ΔminD cells were made chemically competent before transformation with PAR_Amp*. The strains were cultured in LB medium with 50 μg·mL^-1^ kanamycin in a 37 °C incubator.

### Gene circuit design

The designs of both the Positive autoregulation system (Figure S9A) and Simple regulation system (Figure S9B) plasmids used identical modular components (Psal Promoter, GFP, Kanamycin resistance and ColE1 origin of replication).

The codon sequence of salR was designed to fit *E. coli* using GeneOptimizer (ThermoFisher Scientific Ltd.). The salR binding domain and Psal promoter was determined by analysing the sequences of *sal* operon (the GenBank accession no. AF150928). SRS and PRS systems were designed and their structures ware shown in Figure 2A and 2B. In the PAR system, the GFP gene was fused directly to the Psal promoter to replace the sequence of original salA, codon-optimised salR replaced the original salR, and the original structure, promoter and ribosome binding site were kept intact in *sal* operon. In the SRS system, gene optimised salR is under control of a constitutive promoter *pro*D (21), whose structure contained the following elements: promoter-TACTAGAG-B0032-TACTAG-ORF-TACTAGAG-B0015, where the ORF was the salR gene, and B0032 and B0015 are standard Ribosome Binding Sites and terminators from Registry of Standard Biological Parts (www.partsregistry.org). The full sequences of SRS and PAR were synthesised and supplied by GeneArt Gene Synthesis (ThermoFisher Scientific Ltd.). Further detailed information of gene circuit design can be found in Supplementary Information Section 1.

### Culture and Purification of SimCells

Due to the selection marker on *E. coli* MC1000 ΔminD (Kanamycin resistnace), a plasmid of PAR (Psal_GFP_salR) with alternative ampicillin resistance (PAR_Amp*) was needed, and was constructed as follows. The ampicillin resistance gene was isolated by PCR from the plasmid pGEM-T (Promega, UK) using In-Fusion cloning kit (CloneTech, UK) and the following primers: Forward 5’-TTATTGATTGCGGCCTTACCAATGCTTAATCAGTGAGGCACC-3’ and reverse 5’-CACGCCCAGACGGCCCGCGGAACCCCTATTTGTTTATTTTTCT-3’. The resulting sticky end PCR product contained an overhanging EagI site, was thereby ligated in a linearized plasmid PAR. The targeting plasmid PAR was linearized and digested with EagI-HF (New England Biolab), which disrupted the Kan ORF: Only upon correct insertion was ampicillin resistance replaced.

Following heat shock, transformants of *E. coli* MC1000 ΔminD with plasmids PAR_Amp* were screened on Luria-Bertani (LB) agar selection plates containing 100 μg/ml ampicillin. A single colony from each plate was cultured at 37 °C with continuous shaking at 200 rpm overnight in 5 ml LB broth supplemented with 100 μg/ml ampicillin (LB amp). Overnight culture was added to fresh LB amp in a ratio of 1:1000 and cultured for 24 hours. For the minicell purification, a modification of a previously described method (20) was used in order to obtain a high yield and purity, while maintaining maximum ATP within the minicells. Overnight culture was centrifuged at 4 °C at increasing speeds of 1000 g increments, from 1000 g to 4000 g for 10 minutes at each step to remove parent cells from the suspension. The supernatant was subsequently treated with 100 μg/ml ceftriaxone and incubated 37 °C for 1 hr with shaking at 200 rpm, then stored at 4 °C overnight. Following further centrifugation at 4000 g for 15 minutes to pellet any remaining lysed and elongated parent cells as a result of the ceftriaxone treatment, the supernatant was passed through a 0.22 μm nitrocellulose membrane (Sigma, UK) and the minicells re-suspended from the membrane into sterile PBS solution to concentrate them 100-fold. PBS was chosen as it is a cost-effective medium in which to suspend the cells and to maintain osmotic balance. Minicell samples were maintained at 4 °C prior to testing. The purity of minicell suspensions was determined by a plate-count method after 24 h incubation on LB-amp agar plates at 37 °C and conducted in triplicate.

### Flow cytometry

Transformed *E.coli* DH5α cells were induced with gradient of aspirin concentration (final concentrations of 0,0.05,0.1,0.25,0.5,1,2.5,5,10,20,50 μM) in kanamycin (50 μM) LB for 16hrs at in a 600RPM shaker at 37°C. Induced culture was then analyzed using FACSCalibur^™^ (BD Bioscience), with 100,000 events being captured for each sample. Gating was performed on forward, side scatter and GFP channel to remove noise caused by debris and background. Data was exported to Kaluza Flow cytometry for subsequent analysis and visualization.

### Induction and GFP measurements

Stock solution of L-arabinose, glycerol and aspirin was made in deionized water and sterilized using a 0.2μm syringe filter to be tested with cells containing SRS and PAR respectively. A total volume of 200 μL (cultured bacteria cells and inducing aspirin) was loaded in a 96 well black-sided, clear-bottomed plate (Nunclon,UK) in quadruplicate. The plate was then placed into a BioTek Synergy HT microplate reader (BioTek Corporation, UK) maintained at 37°C with reads of fluorescence (excitation:480 nm, emission:520 nm) and optical density (OD) at 600 nm. Readings were recorded every 15 mins for 16 hrs.

For experimental conditions in PBS (phosphate buffer saline) and M9, prior to induction transformed bacteria which were cultured overnight in LB were centrifuged and washed with PBS and M9, respectively, three times at 4°C to remove any residue of LB broth. For achieving micro-aerobic condition, 50 μL of mineral oil was added on top of 200 μL reaction volume in 96 well plate.

### Visualization of bacterial cells

Fluorescence of induced bacterial cells or SimCells was visualized using a Motica BA210 digital microscope with Moticam 580INT display output after 16hrs of induction. 5 μL of each sample was taken from the well and placed on a glass slide with coverslip. Images were taken at 400X magnification. Light micrographs were subsequently analyzed using ImageJ version 1.50b, with fluorescent maxima automatically counted after consistent thresholding.

## Results

### Heterogeneity expression of PAR and SRS circuits in E. coli DH5a

In this study, we initially optimised condons of the salR gene and designed two gene circuits: simple regulation system (SRS) and positive autoregulation system (PAR) and expressed them in *E. coli* DH5a (Fig. 1). In the SRS circuit expression of salR is under the control of a strong constitutive promoter *proD*, due to its good insulation property within different genetic contexts (21). In the PAR circuit salR is controlled by its own promoter to form a fine-tuned feedback regulation loop. *E. coli* DH5a carrying SRS and PAR gene circuits were induced by 0, 2.5 and 50 μM aspirin, and the GFP expression was measured at a single-cell level using flow cytometry and a fluorescent microscope (Fig. 1). SRS has lower background leakiness than PAR. GFP expression in both SRS and PAR can be significantly induced by 2.5 μM aspirin (Fig. 1). Density plots of front scatter reads against GFP channel show a single population of GFP expressing cells in SRS at each inducing concentration, ranging from 3.67 to 32.11 GFP RFU (relative fluorescent units) (Fig.1). The baseline reading in the PAR system was slightly higher (by 7.84 GFP RFU) than that of the SRS system (3.67 RFU). Furthermore, the PAR system exhibits a slightly bistable distribution (Figure 1 and Fig. S1), showing low and high two populations of GFP expressing cells, ranging from 9.69 (Low) to 165.16 GFP RFU (High) (Fig. 1). The leaky and bistable expression of GFP in the PAR system are consistent with classical mathematic models of positive autoregulation circuits (22), confirming that our synthetic PAR and SRS circuits yield behaviours that reflect their original designs.

**Figure 1.**
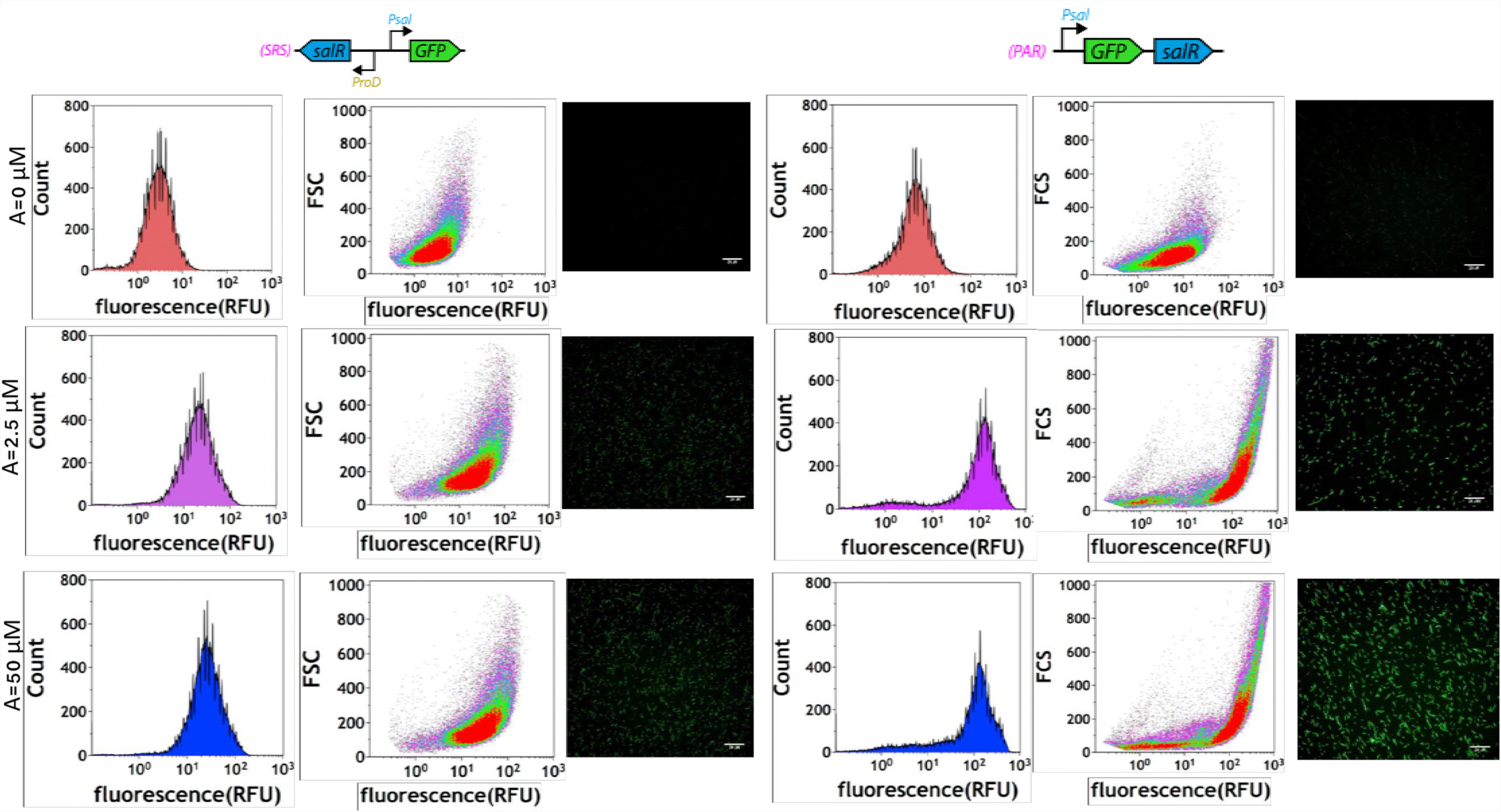
The behavior of single cells in response to aspirin induction was evaluated by flow cytometry. 100,000 cells from un-induced (red), partially induced (magenta) and fully induced (blue) populations were observed for each SRS and PAR circuit configuration. The system configurations were illustrated at top. The specific concentration of aspirin is indicated in the plot. Histograms are plotted with bi-exponential scale to render the wide range of biosensor activation. Together with density plot using FSC(front scattered light) on y-axis and relative fluorescence on x-axis, the presence of large, well-separated bimodal distributions validates that near the bifurcation point, the designed PAR circuit does indeed reflect the ON/OFF induction behavior of a typical positive feedback system.

### Inactivated SalR is able to bind on P_sal_ promoter to repress transcription

The aspirin inducible regulatory system is controlled by the SalR regulatory protein, originally from *Acinetobacter baylyi* ADP1(18). SalR is a LysR-type transcriptional regulator (LTTR) super-family - the largest family of regulators in prokaryotes (23), which contains an N-end DNA binding domain (DBD) and a C-end chemical recognition domain (CRD) (24). Based on previous reports of characteristic features in LTTRs family (25), the “sliding dimer” mechanisms was adopted and illustrated in Fig. S2. This P_sal_-SalR regulation was futher validated using a salicylate biosensor ADPWH_*lux* that was previously constructed by inserting a promoterless *luxCDABE* cassette between *salA* and *salR* of the *sal* operon in *A. baylyi* ADP1 (18, 25).

To validate the molecular mechanism, two SalR mutants ADPWH_NΔsalR and ADPWH_CΔsalR were constructed with N-end and C-end disrupted respectively from ADPWH_*lux* (Table 1). In the mutant ADPWH_CΔsalR, four bases (ATAA) in CDC domain of SalR were deleted using the sacB-km counter selection method (see Supplementary Section 1). We hypothesised that this deletion did not prevent the mutated SalR from binding the DNA at the Psal promoter site, because the N terminus was unaltered. In the case of ADPWH_NΔsalR, the DBD of SalR in ADPWH_KMΔsalR was knocked out by inserting a kanamycin (Km) resistant gene (26) into the ClaI site of SalR in ADPWH_lux. This mutant completely disrupted the DNA binding HTH motif of SalR.

Although both SalR mutants ADPWH_NΔsalR and ADPWH_CΔsalR were unable to respond to salicylate acid, the levels of bioluminescence expression were different (Fig. S3): the average background bioluminescence expression of ADPWH_NΔsalR was 5968±48, which is significantly higher than ADPWH_CΔsalR (3222±66, p< 0.001) and ADPWH_lux without induction (3857±99, p< 0.001). This suggests that the DBD disruption of SalR in ADPWH_NΔsalR could produce a slightly higher bioluminescence due to NΔsalR’s inability to bind the P_sal_ promoter. Since the CRD deletion of SalR in ADPWH_CΔsalR still produced the DNA binding domain, it led the mutant CΔsalR to bind the P_sal_ promoter and repressed the background expression of ADPWH_CΔsalR to the same level as ADPWH_*lux* with an intact SalR (Fig. S3). Thus, the results suggest that SalR occupies P_sal_ promoter regardless of the presence of its inducer.

### SalR_i_/SalR_a_ competitive binding hypothesis and mathematic model construction

Accorrding to the salR mutagenesis experiments above, we hypothese that both inactive SalR_i_ and activated SalR_a_ can competitively bind the P_sal_ promoter. To assess the hypothesised mechanisms and investigate how variation of different aspects of the SRS and PAR systems may affect the aspirin-regulated performance, we developed a mathematical model to simulate and predict the performance of the three gene circuits. The interaction considered in our model for the SRS and PAR systems is outlined in Fig. 2, and is implemented using deterministic differential equations and binding models which describe gene regulation and expression. This process treats the SalR regulatory mechanism as a biased competition between binding of its inducer-bound active form SalR_a_ which activates gene expression; and inactive form SalR_i_ which represses gene expression. For the SRS system, the competition between strong expression of abundant SalR_i_ and aspirin activated SalR_a_ is expected to produce low GFP expression levels. This competition has little effect on the PAR system as its SalR level is tuned by a self-regulated feedback loop.

**Figure 2.**
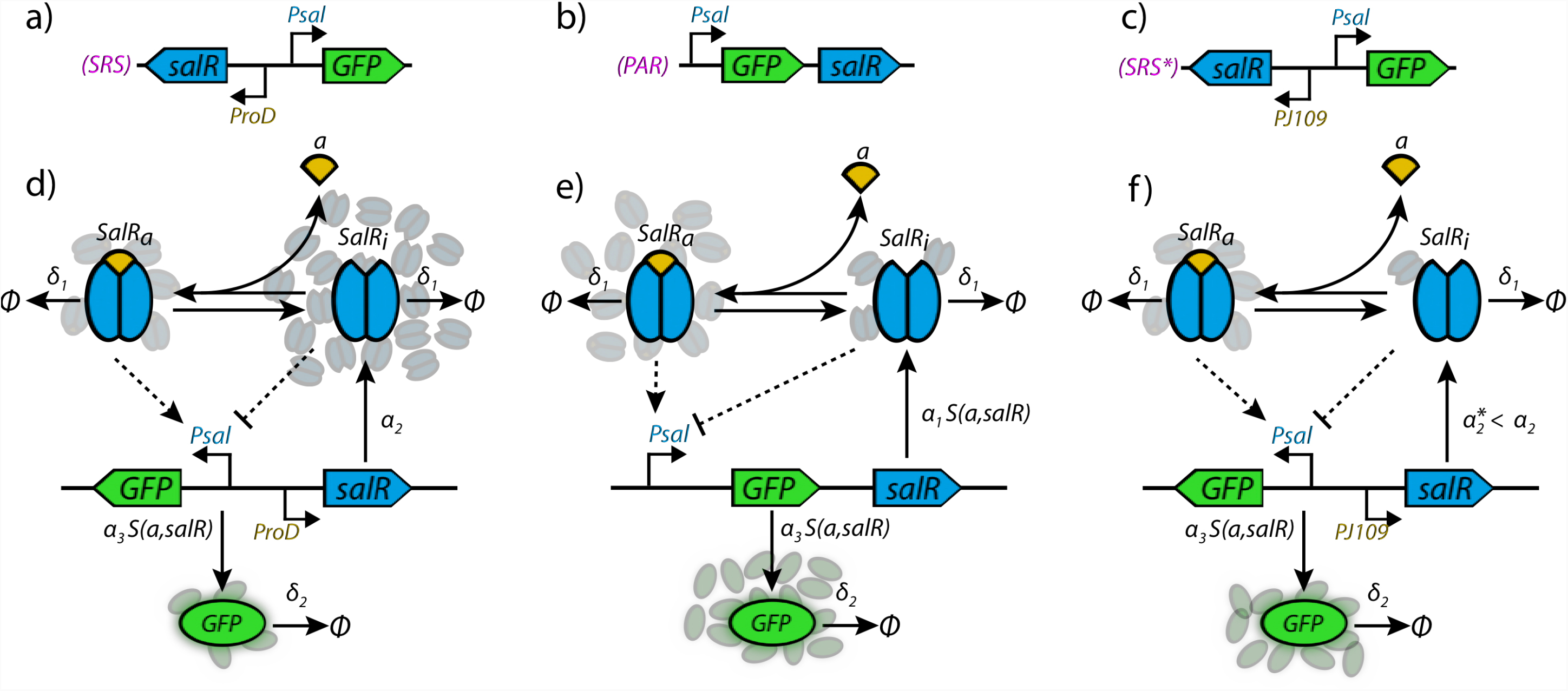
SRS, SRS* and PAR systems and model structure. a) The simple regulatory system (SRS) provides open-loop control of GFP expression by expressing salR from a constitutive promoter (for which we chose the strong promoter proD), which when bound to its inducer activates expression of GFP from regulated promoter Psal. b) The positive autoregulatory system (PAR) closes the loop by expressing both GFP and salR from Psal. Due to transcriptional leakage, a small amount of salR is expressed prior to inducer addition; when inducer is added transcription from Psal is activated, leading to strong expression of both GFP and salR. c) The simple regulatory system (SRS*) variants which strong promoter proD was replaced by a weak promoter PJ109, it still provides open-loop control of GFP expression by expressing salR from a constitutive promoter. e) Model structure for the PAR system. d) Model structure for SRS system and f)odel structure for SRS*. Solid arrows indicate reactions in which the starting species is consumed, whilst dashed arrows mean that consumption is not considered. The extracellular resources (R) and metabolic pathway (P, red box) provide necessary resources for protein expression, producing GFP (green) and salR (blue). salR is assumed to dimerise with high affinity, meaning all is found in its dimer form. An inducer molecule (A, yellow) is transported into the cell to give internal concentration Ai, and then reversibly binds with inactive salR dimer (salRi) to yield its active form (salRa). Two salRa dimers can cooperatively bind Psal to activate gene expression, whilst cooperative binding of two salRi dimers can repress gene expression. For detailed model description and definition of parameters see Supplementary Information Section 1.

We created a two-state differential equation model of the system, in which the concentration of SalR is governed by:

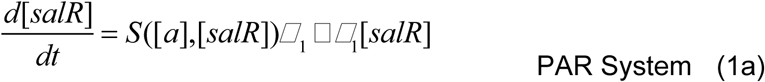

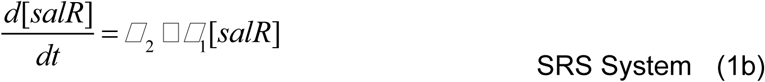

and the system output (GFP) concentration is governed by (for both systems):

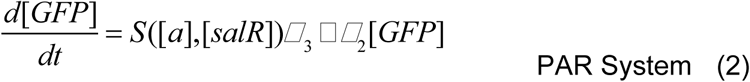

where the regulating function that describes the interactions between salR and inducer concentrations, and the P_sal_ promoter, is given by:

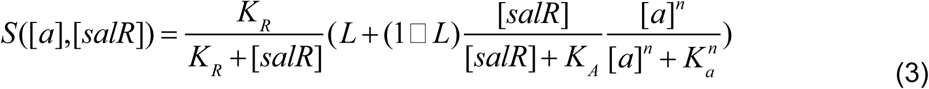

in which *a* is inducer concentration, *L* is the leakage rate from the Psal promoter, *K*_*R*_ defines the saturation point for repressive binding with P_sal_, *K*_*A*_ defines the saturation point for activating SalR binding with P_sal_, and *K*_*A*_ defines the saturation point for inducer interaction with the system. Equation 3 thus reflects the balance we expect between repressive SalR_i_ binding, and inducer-dependent activating SalR_a_ binding. Further discussion of the modelling approach, as well as the procedure used for parameter fitting, is provided in Supplementary Section 2.

### Experimental characterisation and model validation of in E. coli DH5a

The performances of SRS and PAR designs in *E. coli* were characterised in terms of four features: 1) expression level, 2) response time, 3) output dynamic range/induction fold, and 4) input dynamic range and leakiness (un-induced expression level). These four features are motivated by a desire to quantify a biosensor’s utility in drug delivery and *in vivo* sensing. In such an application, a biosensor would ideally exhibit a high sensitivity, fast response, high induction fold change, and tightly controlled expression. To examine these criteria, we ran plate reader experiments to analyse the P_sal_-SalR systems over a range of inducer concentrations, and compared these data to the simulated results from our mathematical model.

The SRS and PAR systems in *E. coli* DH5a were induced by different concentrations of aspirin and the response time of GFP expression was examined in three different growth media (LB, PBS with arabinose, and PBS with glycerol) (Fig. 3, S4 and S5). The resulting expression profiles represent a broad range of sensor behaviours that might arise due to different external disturbances and internal cell-state variation. We defined the time required for biosensor induction response in each condition as the time taken for GFP fluorescence to reach and exceed 10% of its final maximum fluorescence output.

**Figure 3.**
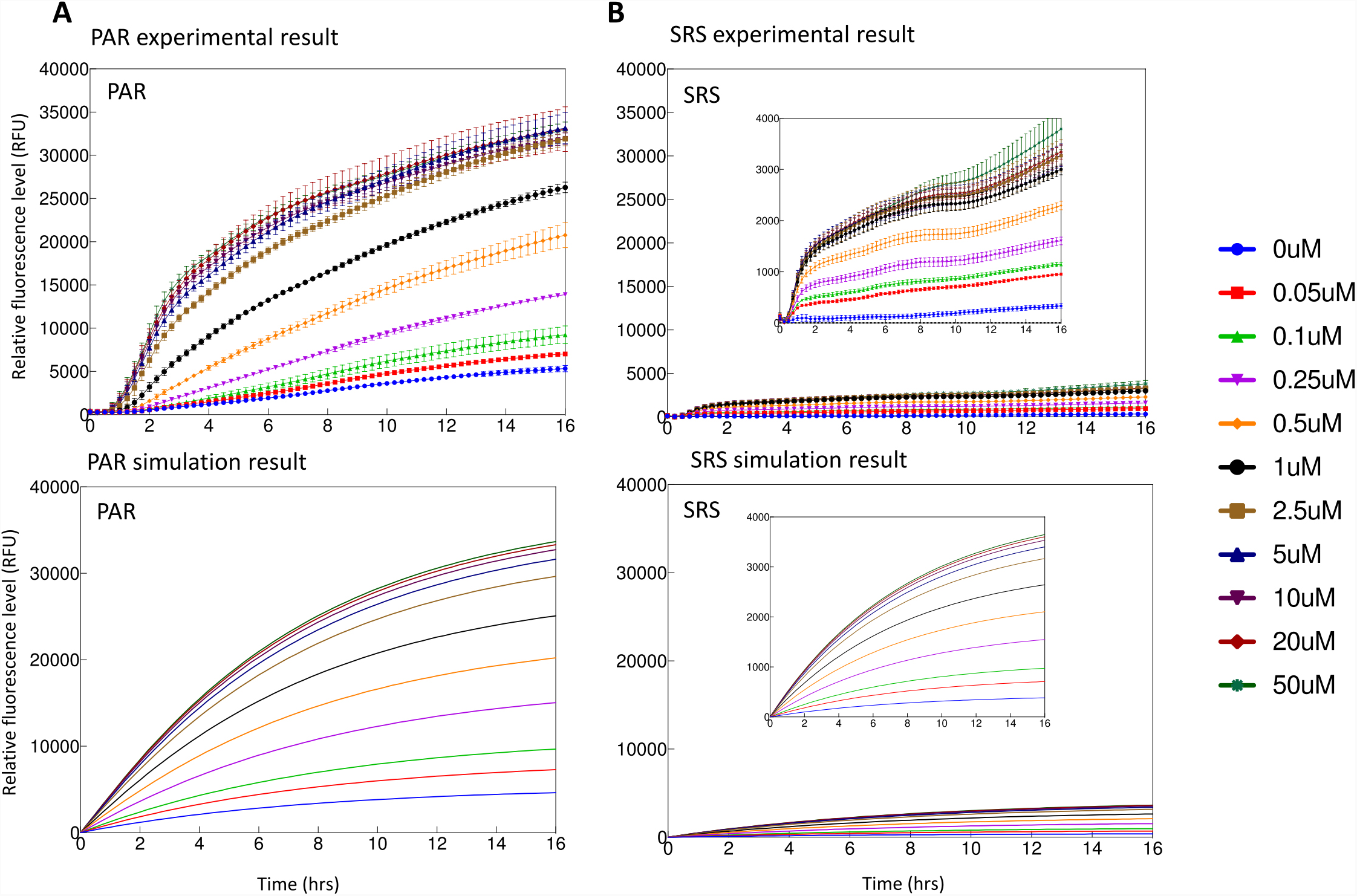
Induction kinetics for the SRS and PAR configurations (A, B) in LB *and in silico* modelling predicts the circuits behaviour in two different design (C,D). Aspirin was added at time zero and fluorescence is observed for 16hrs. Colour coded induction levels are indicated on the right legend. The induction gradient is 0, 0.05, 0.1, 0.25, 0.5, 1, 2.5, 5, 10, 20, 50 μM. Cell intrinsic fluorescence and background was removed. To be easily compared with the experimental data, the simulations are plot with the same inducer concentrations as used for characterization. Standard deviation was plotted for all data replicates (n=4)

The expression profile for both the SRS and PAR circuits in LB media is shown in Fig. 3A and 3B. The fluorescence and OD were monitored for 16 hr over a range of aspirin concentrations between 0.05 to 50 μM. GFP expression in both SRS and PAR circuits was detectable within 50 mins, which is close to the theoretical minimum time required for GFP maturation (27). No significant difference in growth rate was observed between the SRS and PAR systems (Fig. S6). Experimental results from the expression of PAR and SRS circuits are in good agreement with the mathematical model simulation (Fig. 3). Gene expression with the PAR circuit exhibits a substantially higher GFP expression than with SRS circuits. This can be explained as a consequence of the competitive binding of active and inactive SalR protein to P_sal_ promoter (Fig. 2). In SRS system, the constant overexpression of SalRi competitively occupied the P_sal_ promoter, reducing the possibility of active SalR_a_ binding, which repressed overall GFP expression in the presence of aspirin and lowered down the background baseline. The threshold concentration of aspirin required to activate GFP expression in SRS system was 2.5 μM and leaky expression was low (Fig. 3). In contrast, the PAR system tuned the strength of SalR expression through a positive feedback loop and reduces the competitive repression, resulting in much stronger GFP expression with lower activation threshold concentration (0.05 μM) (Fig. 3). However, the level of leaky expression of GFP in PAR system was higher than in SRS circuit.

### Redesign SRS* genetic circuit validate competitive binding hypothesis

To redesign the SRS circuit, the expression of SalR under the control of constitutive promoter with different strength was examined. Promoter *proD* along with three weaker promoters J109, J115 and J106 were fused with the gfp gene separately, and the strenghth was evaluated by detecting GFP expression level. Fig. 4A shows that *proD* was the strongest promoter and J109 was the weakest promoter amongh these four promoters: *proD* J109, J115 and J106, which is consistent to previous reports (21, 28). Subsequently J109 promoter (5 times weaker than *proD*) was characterised and replaced *proD* in constructing SRS* (Fig. 4B). Interestingly, GFP expression level in the SRS* circuit was 3.5 folds higher than original SRS circuit, and the new SRS* showed an intermediate leakniess of GFP expression compared to SRS system (Fig. 3B and 4C). The threshold concentration for SRS* circuit remained the same: 0.05 μM (p< 0.05). This is in a good agreement with the hypothesis, which suggests that the SRS* circuit with the weak promoter J109 should produce less SalR than the SRS circuit, ease the SalR_i_/SalR_a_ competitive binding on P_sal_ promoter and increase GFP expression level (Fig. 4C). By applying the same parameters, our mathematical model fit the experimental results well, and predicts that the transcriptional rate from J109 is ∼15% of that from ProD, similar to the ratio in outputs observed in Fig. 4A. The salR mutagenesis experiment, this experimental results and mathematic model fitting collectively suggest that the hypothesis of SalR_i_/SalR_a_ competitive binding is reasonable.

**Figure 4.**
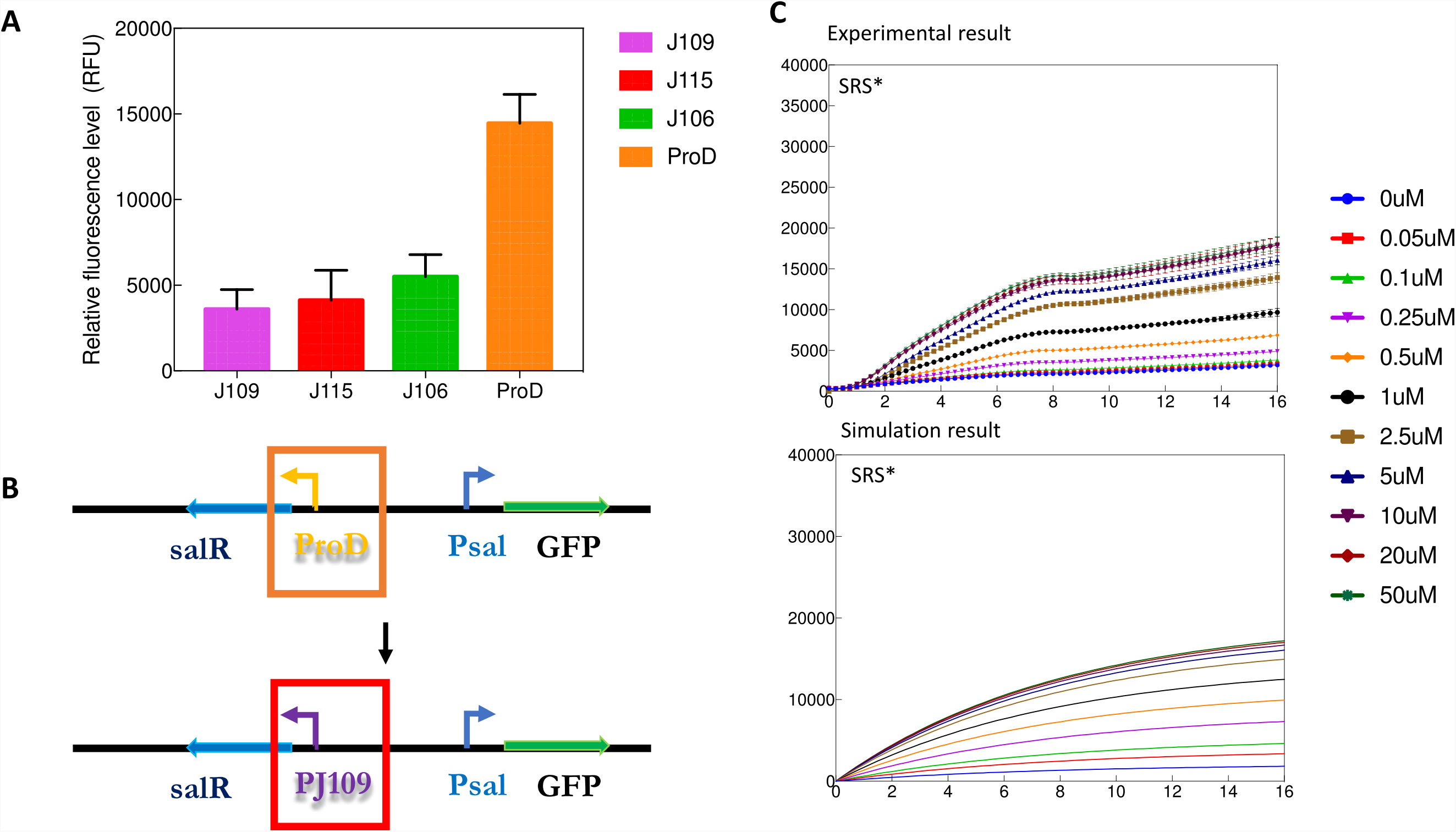
Induction kinetics for the SRS * configurations in LB *and in silico* modelling predicts the circuits behaviour in two different design. A) Schematic diagram of promoter replacement and the configuration of SRS*. B) Aspirin was added at time zero and fluorescence is observed for 16hrs. Colour coded induction levels are indicated on the right legend. The induction gradient is 0, 0.05, 0.1, 0.25, 0.5, 1, 2.5, 5, 10, 20, 50 μM. Cell intrinsic fluorescence and background was removed. To be easily compared with the experimental data, the simulations are plot with the same inducer concentrations as used for characterization. Standard deviation was plotted for all data replicates (n=3) C) The relative fluorescence level was characterised between J109-GFP and ProD-GFP in DH5α.

### Dynamic range and leakiness

The dynamic range of an inducer, also referred to as its *dosage response,* denotes the difference between the induction level that results in the biosensor’s maximal output, and the induction level at which the biosensor’s output increases significantly above the un-induced level. There are a range of metrics for quantifying this criterion, such as the fold-change of inducer concentration over which the sensor goes from 10% of its maximum induction level to 90% of this level (29). Fig. 5 shows that the threshold of SRS, SRS* and PAR activation is 0.05 μM aspirin, GFP expression increases in the range of 0.05-10 μM, and saturates when aspirin concentration is greater than 10 μM. Our mathematical model fits well to the experimental measurements (Fig. 5) in this regard. The corresponding fold-change of inducer concentrations are presented in Table 2. In the LB media experiments, SRS exhibited an induction range from 431±33 to 5,200±650 GFP au and the SRS* showed an intermediate range from 3,201±45 to 18,119±794 GFP.au, whilst PAR exhibited an induction range from 5,325±362 to 32,908±939 GFP.au. (Fig. 5). Notably, PAR had a much higher induction fold change compared with SRS in all carbon condition cases; this in line with what we expected from classical models of autoregulation, which in many established *in-silico* and *in-vivo* studies has demonstrated that one of the benefits of using a positive regulation system architecture is signal amplification (30-32).

**Table 2.** The range of aspirin concentrations over which the system changes from 10% to 90% of maximum induction, and the corresponding fold-change (f.c).

| **Medium** | **SRS system (μM)** | **PAR system (μM)** | **SRS\* system (μM)** |
| --- | --- | --- | --- |
| **LB medium** | 0.1 ~ 2.5 (f.c = 25) | 0.1 ~ 2.5 (f.c = 25) | 0.1 ~ 2.5 (f.c = 25) |
| **PBS+0.02% v/v L-arabinose media** | 1 ~ 10 (f.c = 10) | 1 ~ 10 (f.c = 10) | 1 ~ 10 (f.c = 10) |
| **PBS+0.05% v/v glycerol media** | 1 ~ 10 (f.c = 10) | 1 ~ 10 (f.c = 10) | 1 ~ 10 (f.c = 10) |

**Figure 5.**
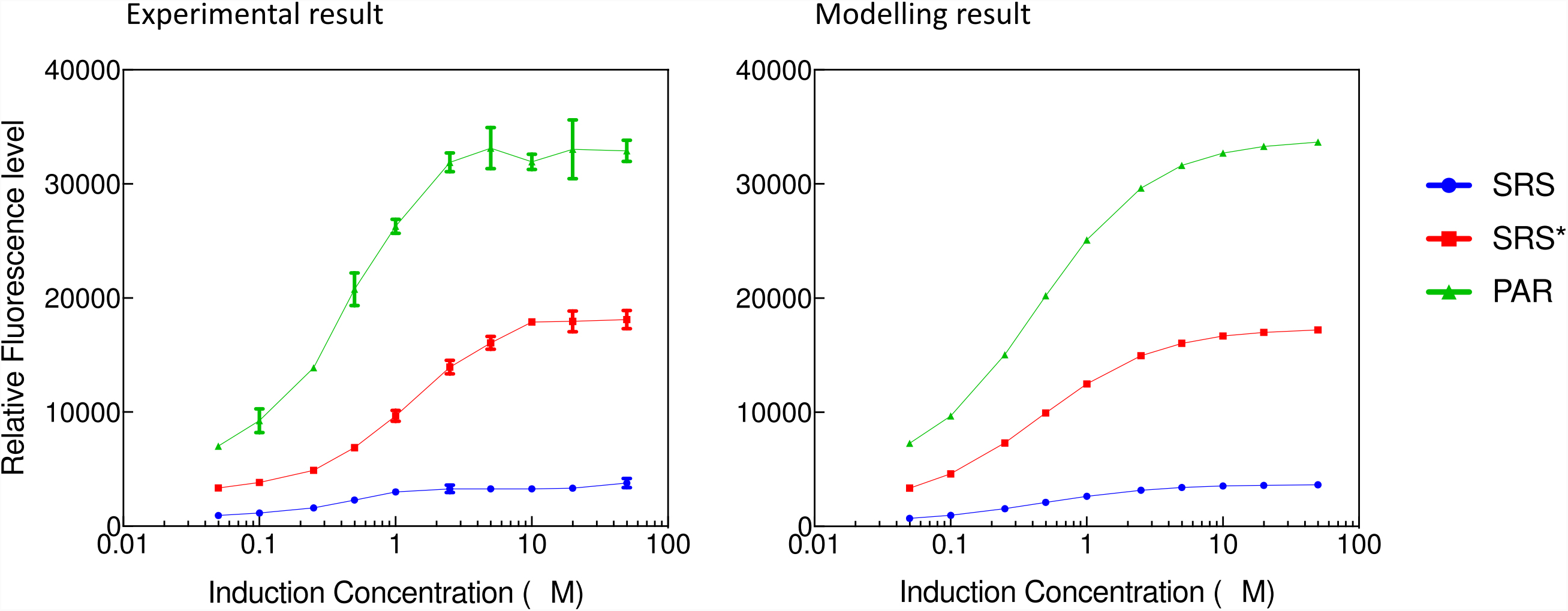
Dosage response curve of the SRS, SRS * PAR configurations in LB. The maximum expression point was determined at the end of a 16hrs induction experiment. The induction gradient is 0, 0.05, 0.1, 0.25, 0.5, 1, 2.5, 5, 10, 20, 50 μM. Cell intrinsic fluorescence and background was removed. The Left diagram corresponds to *in vivo* data, and the right is the corresponding simulation result for each condition. Standard deviation was plotted for all data replicates (n=3).

These results also confirm our conceptual model of the systems considered (Fig. 3), indicating that the SRS system with strong promoter of salR is tightly controlled and has little leaky expression when compared to the PAR system (Table 2). In all experiments of the SRS system, GFP expression levels in uninduced controls and blank background have no detectable difference, and no measurable leakiness was observed over 16 hr, suggesting a very tight control in this design. In contrast, although the PAR system has a strong gene expression, it showed a leaky background particularly when in LB media (Fig. 3). The SRS* increased the expression level from 5,200±650 in SRS to 18,119±794, at a cost of increased leakniess from 431±33 to 3,201±45 GFP.au.

### Performances of SRA and PAR systems in probiotic strain EcN (E.coli Nissle 1917)

The SRS, SRS* and PAR systems were then cloned into a probiotic strain *E. coli* Nissle 1917 (EcN). Considering that the gut environment is semi-aerobic, EcN with both SRS, SRS* and PAR systems was characterised under both aerobic and semi-aerobic conditions. To simulate the potential growth competition with host gut microbiome (33), a nutrient strigented environment was also tested. A small amount of glucose (0.4% w/v) was supplied as sole carbon source, as it is a common carbonhydrate soucre from human diet. EcN with SRS, SRS* and PAR systems maintained a gradient response to inducer concentration in aerobic and semi-aerobic conditions, highlighting our system’s modularity and robustness to a changing chassis and strigent condition (Fig. 6, S6 and S7). The SRS system in EcN exhibited tight control but low response to different concentrations of aspirin, whilst the PAR system demonstrated a higher response to aspirin (Fig. 6). The performances of the SRS, SRS* and PAR systems in EcN were similar to those in *E. coli* DH5a, consistent to the hypothesis of SalR_i_/SalR_a_ competitive binding. Notebly, when supplied with a limited amount of glucose 0.4% (w/v) the expression profile of PAR showed less leakniess, implying that our system could function optimally in nutrient strigent or even deplted condition. We suspect this change might be due to unknown host interference, where cells experiencing glucose starvation alter their resource allocation (34). Futhermore, we noticed there is a slight growth advantages for transformed bacteria in case of 50 μM than in controls (0 μM) (Fig. S7). This applied to all three gene circuits, thus we believed there might be a secondary metabolic breakdown of aspirin involved in *EcN* which is not found in literature before.

**Figure 6.**
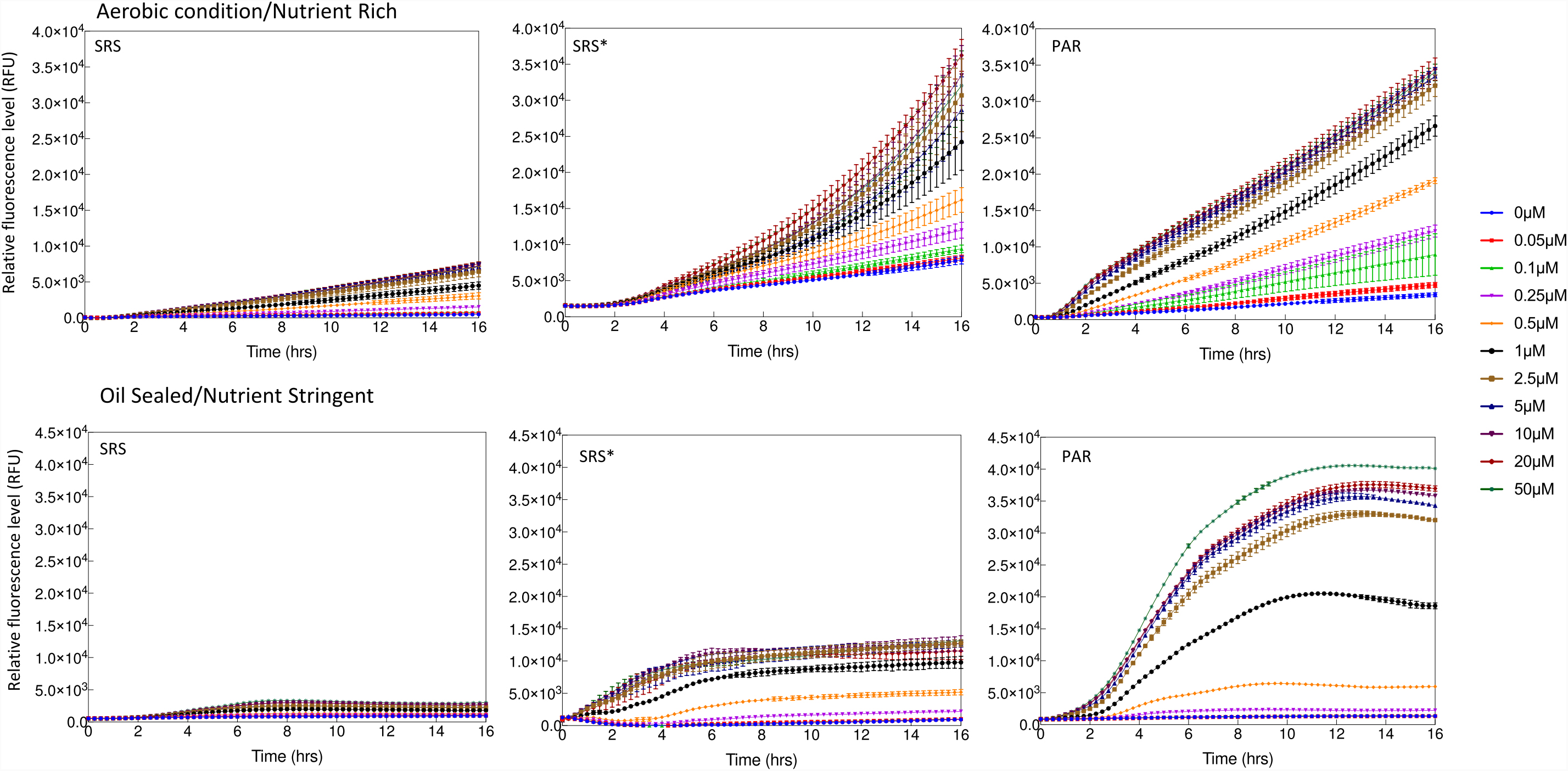
Figure 4. Induction kinetics for the SRS, SRS*, PAR configurations in EcN (*E.coli* Nissle 1917). Aspirin was added at time zero and fluorescence is observed for 16hrs. show the induction kinetics under aerobic condition in LB, Lower panels show the induction kinetics under micro-aerobic condition and nutrient stringent condition (M9 + Glucose 0.4% w/v). Induction levels are colour coded as indicated in the legend. The induction gradient is 0, 0.05, 0.1, 0.25, 0.5, 1, 2.5, 5, 10, 20, 50 μM. Cellular intrinsic fluorescence background was removed.

### Aspirin is converted into salicylate in E. coli

Based on previous work (18), it is evident that the SalR biosensor can be trigged by either salicylate or aspirin. However, it is unclear if aspirin directly activates SalR, or whether its metabolite salicylate does so. Based on the growth advantage we found when cells were supplemented with 50 μM in Figure S7, it is possible that aspirin was metabolised into salicylate and acetate. The later are know to be a food source of *E.coli* in nutrient stringent conditions (35). To investigate the potential mechanism that aspirin could be cleaved by esterase to produce acetate, and salicylate that then activates SalR, we used a Raman probing technique for quick and easy identification.

It has been demonstrated that Raman micro-spectroscopy - deuterium isotope probing (Raman-DIP) can identify metabolically active cells by adding a certain percentage of heavy water (D_2_O) in a medium (36, 37). If cells are metabolically active in the presence of D_2_O, a C-D Raman band (2070 – 2300 cm^-1^) forms via NADPD+ ((H/D free exchange) electon transport chain. As this approach has been demonstrated to be more sensitive compared to tranditional growth curve comparisons, we applied Raman-DIP to explore the activity of the biosensor in response to different carbon sources and thereby the activation pathway (36, 37). Fig. S8A shows single cell Raman spectra (SCRS) of *E. coli* with PAR and SRS systems grown for 16 hr with 35% D_2_O and each with 2 mM glucose, acetate, aspirin or salicylate as the sole carbon source. A clearly distinguishable CD band (2070 – 2300 cm^-1^) was present in the SCRS of cells when they were grown with glucose, acetate and aspirin, indicating cells are metabolically active in these conditions. The CD band was absent when cells were grown with salicylate, since it cannot be metabolised by *E. coli*. As salicylate and acetate are two direct catabolic intermediates from aspirin, our results suggest that *E. coli* should convert aspirin into salicylate and acetate, which drove metabolic activity in cells (Fig. S8). Therefore, although aspirin could not be ruled out as a direct activator of SalR, it is likely that aspirin was broken into salicylate that activates SalR.

### Performance of SRA and PAR systems in SimCells

The SimCells were generated from *E. coli* MC1000 ΔminD with plasmid PAR_Amp* (Table 1). PAR gene circuit was also induced and expressed in purified SimCells (Figure 7 and Figure S9). The lack of any growth by SimCells confirmed their lack of chromosome. Both fluorescent images (Figure S9) and the results from microplate reader (Figure 7) revealed that the background leaky expression of GFP in SimCells was very low, and GFP expression in SimCells became stronger with the increase of aspirin concentrations. SimCells were usually less than 500 nm in size, and some of them were actively moving under microscope (Figure S9), consistent with our previous reports (20).

**Figure 7.**
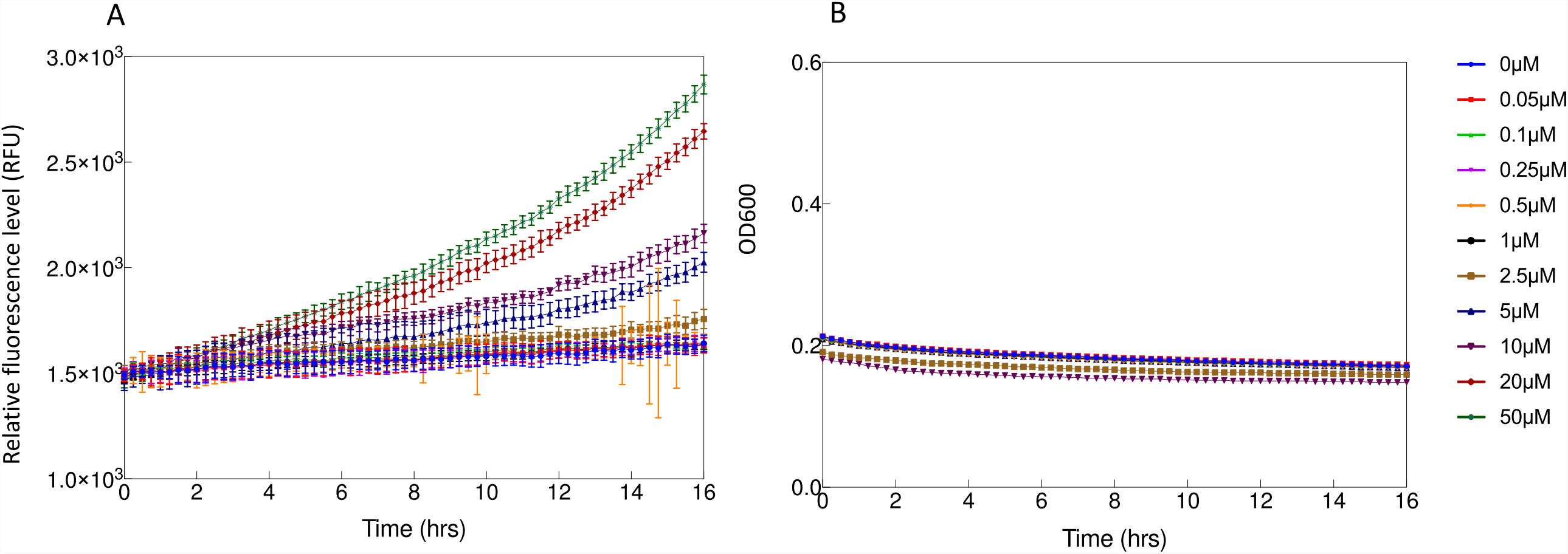
Induction kinetics for PAR* configurations in SimCell. Aspirin was added at time zero and fluorescence is observed for 16hrs. A) show the induction kinetics in PBS medium, B) show the growth kinetics, where no growth was observed and no SimCell was further synthesised. Induction levels are colour coded as indicated in the legend. The induction gradient is 0, 0.05, 0.1, 0.25, 0.5, 1, 2.5, 5, 10, 20, 50 μM. Cellular intrinsic fluorescence background was removed. Standard deviation was plotted for all data replicates (n=3).

## Discussion

### Chassis for P_sal_-SalR system

Probiotic bacteria are safe for human applications, but only a few can be used in gene manipulations (38, 39). *E.coli Nissle* 1917 (EcN) is a safe probiotic bacterium frequently used for medicinal purposes (19, 40-42). Importantly, gene manipulations can be performed in EcN, making it an ideal chassis to synthetic biology. SimCells are small (400-600 nm), chromosome-free and functional cells formed as a result of abnormal cell division (20). SimCells contain transcription and translation machinery, meaning they are able to faithfully express designed gene circuits without interference from background gene networks encoded in the chromosome. Their medical utiity is also aided by the fact that they are non-growing and small in size. Hence, EcN and SimCells were chosen as chassis for the expression of gene circuits in this study.

### A simple and sensitive response to aspirin in P_sal_-SalR system

There is only a limited range of inducers that can effectively operate within the human body, as many inducers may adversely impact human health, or are broadly found in common dietary materials, and should therefore be avoided. For example, antibiotic inducers might alter the structure of microbiota in the gut and sugar inducers may trigger unwanted induction. However, Aspirin is largely absent from common human diets, and its pharmacological safety and effectiveness have been demonstrated extensively through many clinical trials and *in vivo* studies (43, 44). For the SRS and PAR systems studied in this work, the effective range of aspirin dosage required for activation is 0.05-10 μM, which is more than 10 times lower than human safe dosage of aspirin (∼111 μM). Furthermore, induced GFP expression in cells with SRS and PAR systems was observed to start after about 50 min and reached a significant strength within 16 hrs (Figure 3 and 4), which is of a comparable time-scale to Aspirin’s biological half-life.

An aspirin/salicylate triggered NahR regulatory system was previously developed by Royo et al. (9). *Salmonella spp* was engineered for *in vivo* expression of 5-fluorocytosine to treat mice with a fibrosarcoma. This regulatory circuit was a salicylate inducible cascade expression system, which was derived from naphthalene degradative pathway in *Pseudomonas putida* (9). It demonstrated effective drug delivery to minimise tumour size in mice, proving the concept of using salicylate/aspirin as inducers to trigger synthesis of cytosine deaminase that catalysed production of potent cytotoxic fluorouracil for cancer chemotherapy. In this case the gene circuit configuration was a simple cascade expression without feedback, and the effective induction concentration of aspirin was 125-5000 μM (9). Given that aspirin shows toxic effects at concentrations greater than 832 μM (14), sensitive gene circuits for aspirin have an advantage in clinical practice. In addition, the well characterised SRS and PAR systems are simpler gene circuits than regulatory module nahR/Psal::xylS2, demonstrating a dosage response to lower range of aspirin 0.05-10 μM. The importance of feedback loops in genetic circuits is evident in their abundance in naturally occurring gene regulatory systems (22). As a universal network motif in transcription interactions (22, 45), feedback systems can provide several advantages including robustness to noise,(7) improved temporal response(46) and stability in changing cellular contexts (47). The present study not only compared two gene circuits SRS (no feedback) and PAR (with feedback), but also successfully applied a mathematical model to simulate the performance according to LTTR transcriptional regulatory mechanism (Figure 3 and 4). The hypothesised regulatory mechanism was further supported by altering the promoter strength and creating an intermediate SRS* system (no feedback and with a weaker promoter).

The result from our investigation of the molecular mechanism of salR regulation suggest that aspirin/salicylate could be transported into *E. coli* and activate the gene circuits SRS and PAR. In *A. baylyi* ADP1, a putative protein encoded by *salD* is presumably responsible for salicylate transport across the membrane. This protein is homologous to the *E. coli* FadL membrane protein (16). However, it is unclear how efficiently *E. coli* can transport aspirin/salicylate cross the membrane. In *E. coli*, aromatic permeant acids such as salicylate and benzoate induce a large number of low-level multidrug efflux systems, governed by the Mar operon (*marRAB*) as well as additional, unidentified mechanisms (48). Since *E. coli* does not have a specific transport protein for aspirin/salicylate, it is likely the efflux system will remain on after the induction, we hypothesise that there is a maximum equilibrium concentration of aspirin/salicylate inside the cell cytoplasm. This hypothesis works well in mathematic models to successfully simulate the performance of *E. coli* in response to different concentrations of aspirin and growth conditions.

### The performance of P_sal_-SalR system is robust in various conditions

In the present study, many aspects of the salR regulatory system were investigated: its competitive inducer binding properties, response to varying inducer concentrations, and behaviour in *E. coli* given varying system architectures and growth media conditions. To tie the diversity of experimental observations together, and to test the hypotheses that they inform, we built a mathematical model and demonstrated that it was able to describe and simulate all aspects of our systems behaviour. Not only does thes model support the mechanism that we proposed underlies the performance of both the PAR and SRS systems, but it also provides a useful tool for the development of different systems that employ the SalR (or similar) regulatory system. These may include synthetic biological devices designed for clinical applications in the future; in such cases *a priori* utilisation of the modelling architecture developed herein can be used to inform design choices and experimental approaches, thereby optimising designs prior to their implementation.

Since many genera of bacteria naturally co-exist with tumours in clinical situations (2), probiotic bacteria such as EcN and non-growing SimCells should be ideal chasses to construct anti-cancer agents due to their designable features, safe non-proliferation properties, and small size. Controllable expression of gene circuits, combined with safe delivery and tissue penetration would enable EcN and SimCells to be selectively exerting cytotoxic to tumours. This study demonstrates that aspirin inducible SalR system is able to function in EcN and SimCells, which would be useful for bacterial diagnosis and therapy in medicine.

## Acknowledgements

WEH acknowledges support from EPSRC (EP/M002403/1 and EP/N009746/1). AP acknowledges support from EP/M002454/1. HS acknowledges support of the General Sir John Monash Foundation. We also thank Dr Lingjuan Wu for providing strains and scientific advice.

## Supplementary figure caption

**Figure S1**. A) Microscope images of SRS and PAR. Images were taken at both GFP channel and overlaid with Bright field channel for indication of potential mutated circuits or non-transformed bacterial cells. The specific concentration of aspirin is indicated beside the plot. On the left, enlarged images of the PAR circuit when induced with aspirin (50uM) clearly show two separate populations with different GFP intensity (High/Low) as reflected in result from flow cytometry. B) Negative and blank control of aspirin induction experiment in flow cytometry. Cells were sonicated and gated to ensure GFP reads are only from intact & functional cells.

**Figure S2**. A potential schematic for the salR “sliding dimer” regulation mechanism. (A) In the absence of SA, salR regulates the expression of the Psal operon by binding the promoter region at three different functional subsites: a high affinity Repression binding site (RBS), often found near position -65 relative to the transcription start site and two low affinity activation binding sites (ABS’ and ABS’’) found near positions - 10 and -35 respectively. (B) upon activation by aspirin binding, a shift in promoter region binding sites from RBS/ABS’ to RBS/ABS’’ releases the -25 box of the promoter region DNA and leading to (C) RNA polymerase recognition and subsequent gene expression.

**Figure S3**. A) Bioluminescence response to SA of ADPWH_lux, ADPWH_CΔsalR and ADPWH_NΔsalR. ADPWH_lux has a strong, rapid, response to SA, while the two mutants with incomplete salR do not respond at all. Standard deviation was plotted for all data replicates (n=4), while SD for some data sets were insignificant and the enlarged picture was provided in B) as a clear indication of increased background level in case of ADPWH_NΔsalR.

**Figure S4.** The effect of carbon source and growth in designed systems. Induction kinetics for the SRS and PAR configurations in (Top) PBS supplemented with Glycerol (0.05%v/v) and PBS supplemented with 0.02% v/v L-arabinose (Bottom). Aspirin was added at time zero and fluorescence is observed for 16hrs. Lower panels show the *in silico* simulation result. Colour coded induction levels are indicated on the right legend. The induction gradient is 0, 0.05, 0.1, 0.25, 0.5, 1, 2.5, 5, 10, 20, 50 μM. Cell intrinsic fluorescence and background was removed. Standard deviation was plotted for all data replicates (n=4)

**Figure S5**. Growth kinetics for the SRS and PAR configurations PBS supplemented with 0.02% v/v L-arabinose, PBS supplemented with 0.05% v/v glycerol. Top row illustrates that both the SRS and PAR systems have similar growth from OD 0.2 to OD 0.42 over 16hrs. The middle row and bottom row (PBS supplemented with Arabinose and Glycerol respectively) indicate that both strains have no/minor growth over 16hrs. Standard deviation was plotted for all data replicates (n=4).

**Figure S6**. Growth kinetics for the SRS SRS* and PAR configurations in EcN under micro-aerobic condition and LB medium. Standard deviation was plotted for all data replicates (n=3).

**Figure S7**. Growth kinetics for the SRS SRS* and PAR configurations in EcN under micro-aerobic and nutrient stringent condition (M9 medium supplemented with Glucose 0.4%w/v). Standard deviation was plotted for all data replicates (n=3).

**Figure S8**. A) Single-cell Raman spectra of cells fed overnight with 2 μM glucose, acetate, aspirin or salicylate. The C-D band at 2070 – 2300 cm^-1^ is highlighted which demonstrates the general cellular metabolism in the presence of different carbon sources. Each spectrum is an average of 15-25 single-cell Raman spectra and the shaded area represents the standard deviation. B) Box plots of single-cell C-D band integration at 2070 – 2300 cm^-1^ show statistical distribution of D content in SCRS of SRS and PAR in the presence of 2 mM glucose, acetate, aspirin and salicylate, respectively. Significance was compared between salicylate (negative control) and aspirin with P< 0.0001.

**Figure S9**. Microscope images of PAR_Amp* transformed SimCell. Images were taken at both GFP channel and overlaid channel for indication of potential mutated circuits or non-transformed Simcells. The specific concentration of aspirin used is indicated above plot. Background was filtered using ImagJ.

Characterisation and development of aspirin inducible biosensors in *E. coli* Nissle 1917 and SimCells

